# Therapeutic inhibition of monocyte recruitment prevents checkpoint inhibitor-induced hepatitis

**DOI:** 10.1101/2023.08.14.553197

**Authors:** Cathrin LC Gudd, Stephen R Atkinson, Eoin Mitchell, Marie-Anne Mawhin, Samra Turajlic, James Larkin, Mark R Thursz, Robert D Goldin, Nick Powell, Charalambos G Antoniades, Kevin J Woollard, Lucia A Possamai, Evangelos Triantafyllou

## Abstract

Checkpoint inhibitor-induced hepatitis (CPI-hepatitis) is an emerging problem with the widening use of CPIs in cancer immunotherapy. Here, we developed a mouse model to characterise the mechanism of CPI-hepatitis and to therapeutically target key pathways driving this pathology. C57BL/6 wild-type (WT) mice were dosed with TLR9-agonist (TLR9-L) for hepatic priming combined with anti-CTLA-4 plus anti-PD-1 (CPI) or control (PBS) for up to 7 days. Co-administration of CPIs with TLR9-L induced liver pathology closely resembling human disease, with increased infiltration and clustering of granzyme B^+^perforin^+^CD8^+^ T cells and CCR2^+^ monocytes, 7 days post treatment. This was accompanied by apoptotic hepatocytes surrounding these clusters and elevated cytokeratin-18 and alanine transaminase plasma levels. Liver RNA sequencing identified key signalling pathways (JAK-STAT, NF-_κ_B) and cytokine/chemokine networks (*Ifnγ, Cxcl9, Ccl2/Ccr2*) as drivers of CPI-hepatitis. Using this model, we show that CD8^+^ T cells mediate hepatocyte damage in experimental CPI-hepatitis. However, their liver recruitment, clustering, and cytotoxic activity is dependent the presence of CCR2^+^ monocytes. Absence of hepatic monocyte recruitment in Ccr2^rfp/rfp^ mice and CCR2 therapeutic inhibition by cenicriciroc (CVC) in WT mice prevented CPI-hepatitis. In conclusion, using this newly established mouse model, we demonstrate a central role of liver infiltrating CCR2^+^ monocyte interaction with cytotoxic CD8^+^ T cells in the pathogenesis of CPI-hepatitis and highlight novel therapeutic targets.

## Introduction

Checkpoint inhibitors (CPIs) are a class of cancer immunotherapy with proven efficacy in a range of malignancies (1–3). These drugs are monoclonal antibodies targeting immune checkpoint receptors such as cytotoxic T lymphocyte antigen-4 (CTLA-4), programmed cell death-1 (PD-1) and its ligand programmed cell death-ligand-1 (PD-L1) (1). Under normal physiological conditions, immune checkpoints are major regulators of immune homeostasis and tolerance, preventing the generation of auto-reactivity and immune-mediated tissue damage (3). Blockade of these immune checkpoints has been shown to enhance cellular immunity and stimulate anti-tumour responses which result in improvements in long term survival of patients with a range of cancer types (4–7).

As checkpoint molecules are involved in pivotal regulatory pathways, CPI therapy is frequently complicated by immune-related adverse events (irAEs) (8–10). These side effects may manifest in several different tissues (9–11). CPI-induced hepatitis (CPI-hepatitis) is one of the most common organ toxicities affecting up to 30% of patients on dual agent CPI therapy (5,12–15). CPI-hepatitis is associated with increased serum alanine transaminase levels (ALT) levels and characterised histologically by lobular liver inflammation, with CD8^+^ T cell and macrophage inflammatory clusters associated with hepatocyte injury. The severity of disease can range from incidental, mild liver function test abnormalities to fulminant hepatic failure and death (14–19). Current clinical care relies on cessation of CPI therapy and treatment with broadly immunosuppressive treatments, particularly corticosteroids, which have significant off target side effects (20,21). A pressing clinical need therefore exists, to better understand what triggers liver inflammation in a proportion of CPI-treated patients and the immunopathology of this condition to develop targeted treatments.

The liver is constantly exposed to microbial and non-microbial products, including TLR ligands (TLR-L) from the portal blood stream. Thus, robust regulatory mechanisms are required to promote liver tolerance and avoid excessive inflammatory responses in homeostasis (22–25). Immune checkpoint pathways play an essential role in mediating this tolerance (23,26–28).

In the absence of immune checkpoint signalling, either through blockade (e.g. anti-CTLA-4, anti-PD-1) or genetic deletion (e.g. PD-1^-/-^), mice do not spontaneously develop hepatic inflammation (29–31). However, studies utilising mouse models of drug-induced liver injury (DILI) report the induction of more severe and persistent toxicity when hepatic tolerance is broken by interruption of CPI signalling (29,30,32). For example, Metushi et al. reported hepatocyte injury and enhanced effector T cell function in PD-1^-/-^ mice treated with anti-CTLA-4 and Amodiaquine, a known direct liver toxin (29). In addition, Affolter et al. and Llewelyn et al. have shown that administration of anti-CTLA-4 and an IDO1 inhibitor in PD-1^-/-^ mice induces enhanced liver toxicity, reproducing aspects of human CPI-hepatitis (30,33). Evidence from patients treated with sequential anti-cancer agents has suggested that drugs, which are minimally hepatotoxic when administered alone, can trigger significant liver injury in patients with prior checkpoint inhibitor exposure (34,35). This suggests that the inhibition of tolerance pathways by checkpoint blockade, promotes exaggerated immune-mediated liver damage in response to relatively minor direct hepatotoxic triggers.

Here, we demonstrate that TLR9 agonist (TLR9-L) challenge of anti-CTLA-4/anti-PD1 treated mice reproduces key characteristic features of human CPI-hepatitis, including an inflammatory infiltrate of CCR2^+^ monocytes and cytotoxic CD8^+^ T cells associated with surrounding hepatocyte apoptosis. RNA sequencing further confirmed the highly inflammatory environment and highlighted a complex intra-hepatic immune crosstalk. Using Ccr2^rfp/rfp^ transgenic mice in this model, we show that CCR2^+^ liver-infiltrating monocytes are necessary for the recruitment and activation of tissue-destructive CD8^+^ T cells. Furthermore, we demonstrate that therapeutic inhibition of monocyte liver recruitment using cenicriciroc (CVC; CCR2/5 antagonist), prevented the development of CPI-hepatitis.

## Methods

### Animals

All experimental protocols were approved by Imperial College London in accordance with U.K. Home Office regulations (PPL: P8999BD42). C57BL/6 wild-type (WT) male mice (8-12 weeks old) were purchased from Charles River Laboratories. Rag2^-/-^ [B6(Cg)-Rag2t^m1.1Cgn^/J] and Ccr2^rfp/rfp^ [B6.129(Cg)-Ccr2^tm2.1Ifc^/J] mice were kindly donated by Professor Marina Botto and Dr Kevin Woollard, respectively (Imperial College London). All mice were housed and bred under the same specific pathogen-free conditions at the Imperial College London animal facility.

### Animal treatments

Anti-PD-1 (clone RMP1-14), anti-CTLA-4 (clone 9H10), anti-CD4 (clone GK1.5) and anti-CD8 (clone YTS169.4) monoclonal antibodies (mAbs) were purchased from BioXCell, diluted in sterile phosphate buffered saline (PBS) and injected intraperitoneally (i.p.) at a concentration of 200μg/mouse.

Combination anti–PD-1 and anti-CTLA-4 (“CPI”) or sterile PBS as control (“PBS”) were given every two days on day 0 (D0), D2 and D4. On D1, 20 μg/mouse of TLR9-L (CpG oligodeoxynuleotide 1668: 5-S-TCCATGACGTTC CTGATGCT-3) (TIB Molbiol, Germany) was administered i.p. to prime hepatic inflammation. Mice were sacrificed, and blood and liver tissue were collected on D1, D4 and D7 as described in **Supplementary Methods**.

Anti-CD4 or anti-CD8 mAbs were given on D-2 and D-1 before and on D3 after the first CPI injection. 100mg/kg/day of Cenicriviroc (CVC) mesylate (MedchemExpress, USA) or vehicle control was administered in drinking water 4 days before the first CPI injection and for the whole duration of the time course (36). Mice were sacrificed on D7 and blood and liver tissue were harvested.

### Plasma screening

#### Liver Function Tests

Liver function was assessed by measuring alanine transaminase (ALT) (service provided by MRC Harwell Institute, UK) and cytokeratin-18 (CK-18) levels in plasma. CK-18 was measured by enzyme-linked immunosorbent assay (ELISA) (Abcam, UK). The optical density was measured at 450 nm using the Multiskan Go plate reader (Thermo Fisher Scientific, UK).

#### CXCL9 and CXCL10 levels

Levels of CXCL9 and CXCL10 were measured in plasma (diluted 2-fold) using the mouse cytokine release syndrome panel LEGENDplex system (BioLegend, UK), according to manufacturer’s instructions. The acquisition was performed on the BD LSRFortessa™.

### Liver histology and immunofluorescence

Formalin-fixed paraffin-embedded (FFPE) liver sections stained with haematoxylin and eosin (H&E) were provided by the Research Histology Facility, Imperial College London. Immunofluorescent staining of OCT fixed liver cryosections was performed to assess the expression of CD8, F4/80, CD11b, CCR2, granzyme B (GZMB) and albumin using fluorochrome-labelled mAbs listed in **Supplementary Tab.1**. Slides were mounted with fluoroshield with DAPI (Sigma-Aldrich, USA). Images were captured using the Leica DM4 B microscope and the LAS X 3.3.3.16958 software (Leica Camera AG, Germany).

### Quantitative reverse transcription PCR (RT-qPCR)

Total mRNA was extracted from snap frozen liver tissue using the RNeasy Plus Mini Kit (Qiagen, Germany) and reversed transcribed to cDNA using SuperScript IV reverse transcriptase with Random Hexamers (Invitrogen, USA), according to manufacturer’s instructions. The 2xSensiMix SYBR Lo-ROX kit (Bioline, UK) was used for quantification of *Cxcl9*, *Cxcl10* and *Ccl2* mRNA levels (**Supplementary Tab.3**). qPCR was performed using the Applied Biosystems Viia7-fast block instrument and analysed with the Viia7 software (Thermo Fisher Scientific, UK). Fold-change in gene expression was calculated as 2^-ΔΔCt^, as compared to D1 control mice (n=3) of the respective strain.

### RNA sequencing

RNA was extracted from snap frozen liver tissue of WT mice on D1, D4 and D7 (n=4/group) post administration of CPI or PBS, following manufacturer’s instructions using the RNeasy Plus Mini Kit (Qiagen, Germany). RNA sequence data was aligned to GRCm39/Ensembl v104 with transcript quantification by RSEM. Count data were quality controlled and analysed in R v4.0.4 using limma-voom, dorothea and gage. A detailed description of all related methods is provided in supplementary materials.

### Flow cytometry of liver immune cells and absolute cell counts

Surface and intracellular staining of isolated hepatic mononuclear cells were carried out using fluorochrome-labelled mAbs listed in **Supplementary Tab.4**. The acquisition was performed on the BD LSRFortessa™ and data were acquired using the BD FACSDiva™ software (Becton Dickinson Ltd, UK). Data was analysed with FlowJo v10.9.0. Absolute cell counts were obtained using the 123count eBeads (Thermo Fisher Scientific, UK) by flow cytometry, according to the manufacturer’s instructions.

### Statistical analysis

Graph Pad Prism Software 9.5.1 was used to test for statistical significance. Analyses were conducted to compare different groups at one given time point (D1, D4 or D7) or in comparison to D1 within each respective group. For non-parametric un-paired analysis, Mann-Whitney *U* tests were performed comparing two groups and Kruskal-Wallis tests were used for comparisons between three or more groups. For correlations, Spearman rank correlation coefficients were used. Data are expressed as mean ± standard error of mean (SEM) and statistical significance is assumed for p values of less than 0.05.

Other details and additional experimental procedures are provided in the supplementary methods.

## Results

### CTLA-4/PD-1 blockade in combination with TLR9-L administration induces liver inflammation and hepatocyte damage

In line with previous reports (29–32), breaking immune tolerance by treating mice with combination anti-CTLA-4/anti-PD-1 (CPI) was insufficient in inducing hepatitis in mice (**Supplementary Fig.1 A**). Here, we sought to examine whether administration of TLR9-L in WT mice, mimicking the microbial signal (unmethylated CpG rich DNA) from viral/bacterial infection or increased intestinal bacterial translocation into the portal bloodstream, in combination with CPI treatment would induce liver inflammation to model human disease (**Fig.1 A**) (18).

**Figure 1.**
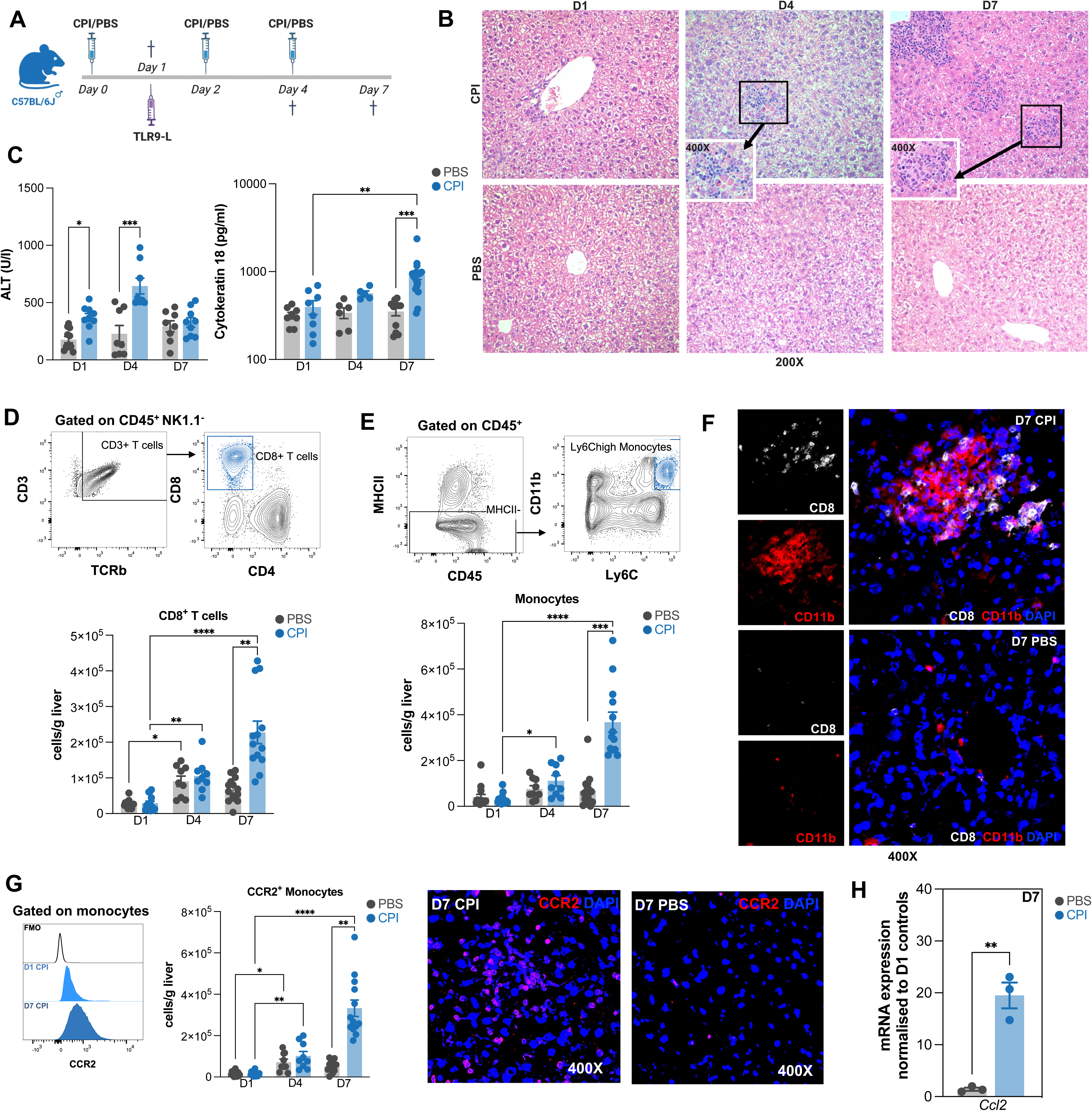
Liver inflammation and hepatocyte damage following TLR9-L/CPI treatment of mice. A) Set-up of experimental CPI-hepatitis time course, comparing CPI (blue) and PBS (grey)-treated mice across different time points. B) Representative H&E stained liver sections of CPI or PBS-treated mice during the time course (n=8/group/time point). Magnification: 200X, 400X. C) Measurement of liver damage indices (ALT, CK-18) in plasma. D) Representative contour plots and absolute numbers of total liver CD8^+^ T cells. E) Representative gating strategy and absolute numbers of total liver monocytes during the time course. F) Representative pictures of liver cryosections stained for CD8 (white), CD11b (red) and DAPI (blue) on D7 of CPI and PBS-treated mice (n=8/group). Magnification: 400X. G) Histograms and absolute number of CCR2^+^ monocytes and representative pictures of liver cryosections stained for CCR2 (red) and DAPI (blue) of CPI and PBS-treated mice on D7 (n=4/group). Magnification: 400X. H) Fold change expression of total mRNA levels of *Ccl2* in liver homogenates on D7, normalised to respective D1 controls (n=3). Each symbol represents an individual mouse. Results shown are representative of 1-3 independent experiments. * p<0.05, ** p<0.01, *** p<0.001. **** p<0.0001.

Following a single dose of CPI or PBS and prior to TLR9-L administration (D1), mice showed normal liver histology (**Fig.1 B**). However, after TLR9-L dosing, CPI-treated mice showed panlobular or centrilobular granulomatous hepatitis on D4 (**Fig.1 B**), with even more pronounced inflammatory foci on D7, similarly to human liver pathology (15,37). Notably, apoptotic and swollen hepatocytes were seen surrounding hepatic immune clusters (**Fig.1 B**).

To assess the severity of liver injury, ALT and CK-18 levels were measured in plasma. Although liver histology was normal on D1 (**Fig.1 B**), CPI-treated mice showed increased ALT levels, compared to PBS-treated mice on D1 (**Fig.1 C**). ALT levels increased further on D4 in CPI-treated mice, but remained stable in PBS-treated mice. In addition, CK-18 was elevated in CPI-treated mice on D7, compared to D7 PBS and D1 CPI (**Fig.1 C**). Of note, hepatic TLR4-L priming induced milder liver inflammation on D7 following CPI treatment compared to TLR9-L priming, but had no effect on ALT levels (**Supplementary Fig.1 B-D**).

### Liver infiltration of CD8^+^ T cells and CCR2^+^ monocytes in TLR9-L/CPI-treated mice

Studies on human CPI-hepatitis from our group and others report an increased recruitment and aggregation of CD8^+^ T cells with CCR2^+^ monocytes in histological observations (15,18,37). These inflammatory structures are associated with surrounding hepatocyte death and are uniquely found in CPI-hepatitis (37). To investigate whether this is recapitulated in our model, we used flow cytometry and immunofluorescence microscopy to identify and quantify the liver myeloid and lymphoid populations (38–41).

No differences in NK, NKT, B, and CD4^+^ T cell populations were observed between CPI and PBS-treated mice (**Supplementary Fig.2 A-C**). However, liver CD8^+^ T cells were significantly increased in CPI-treated animals on D7, compared to PBS (**Fig.1 D**). Similarly, Ly6C^high^ inflammatory monocytes were expanded on D7 compared to control (**Fig.1 E**). In addition, D7 CPI-treated mice showed significantly increased numbers of resident Kupffer cells (KC) and monocyte-derived macrophages (MoMF), compared to D7 PBS (**Supplementary Fig.3 A+B**).

**Figure 2.**
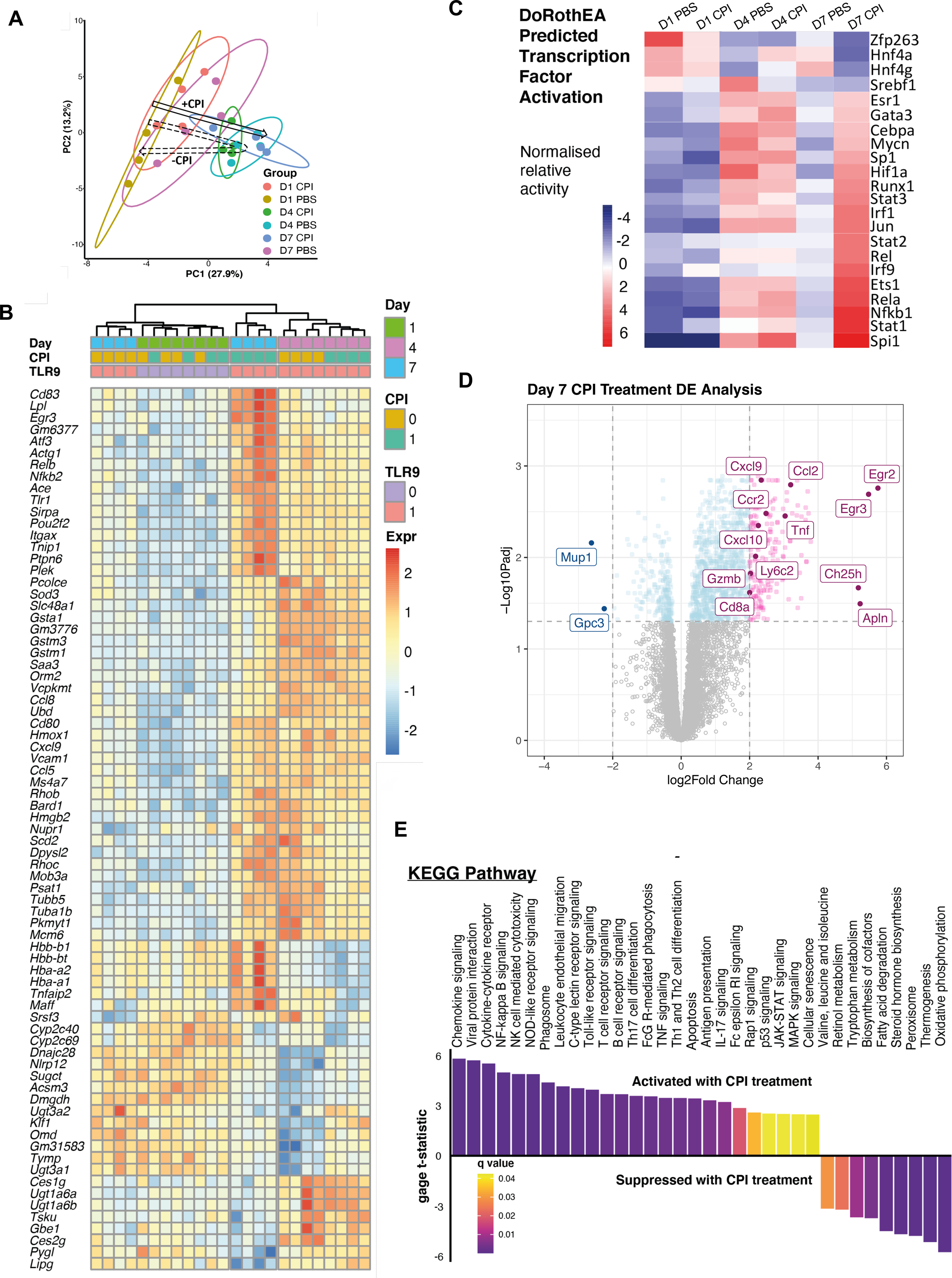
Liver transcriptome demonstrates perpetuated inflammatory responses with combined TLR9-L/CPI treatment. RNA sequencing analysis of n=4 mice/group for D1, D4, D7. A) A schematic summary of principal components analysis (PCA) with a transient shift induced by TLR9-L in the absence of CPI (dashed arrow) but sustained in its presence (solid arrow). B) Heatmap of differentially expressed genes across all samples. C) Activity of selected transcription factors inferred using dorothea and summarised by time point and treatment group. D) Volcano plot of differential expression analysis between mice after 7 days of treatment with, or without, CPI. E) Significantly dysregulated KEGG pathways tested using Generally Applicable Gene set Enrichment analysis for D7 CPI compared to D7 PBS as baseline.

**Figure 3.**
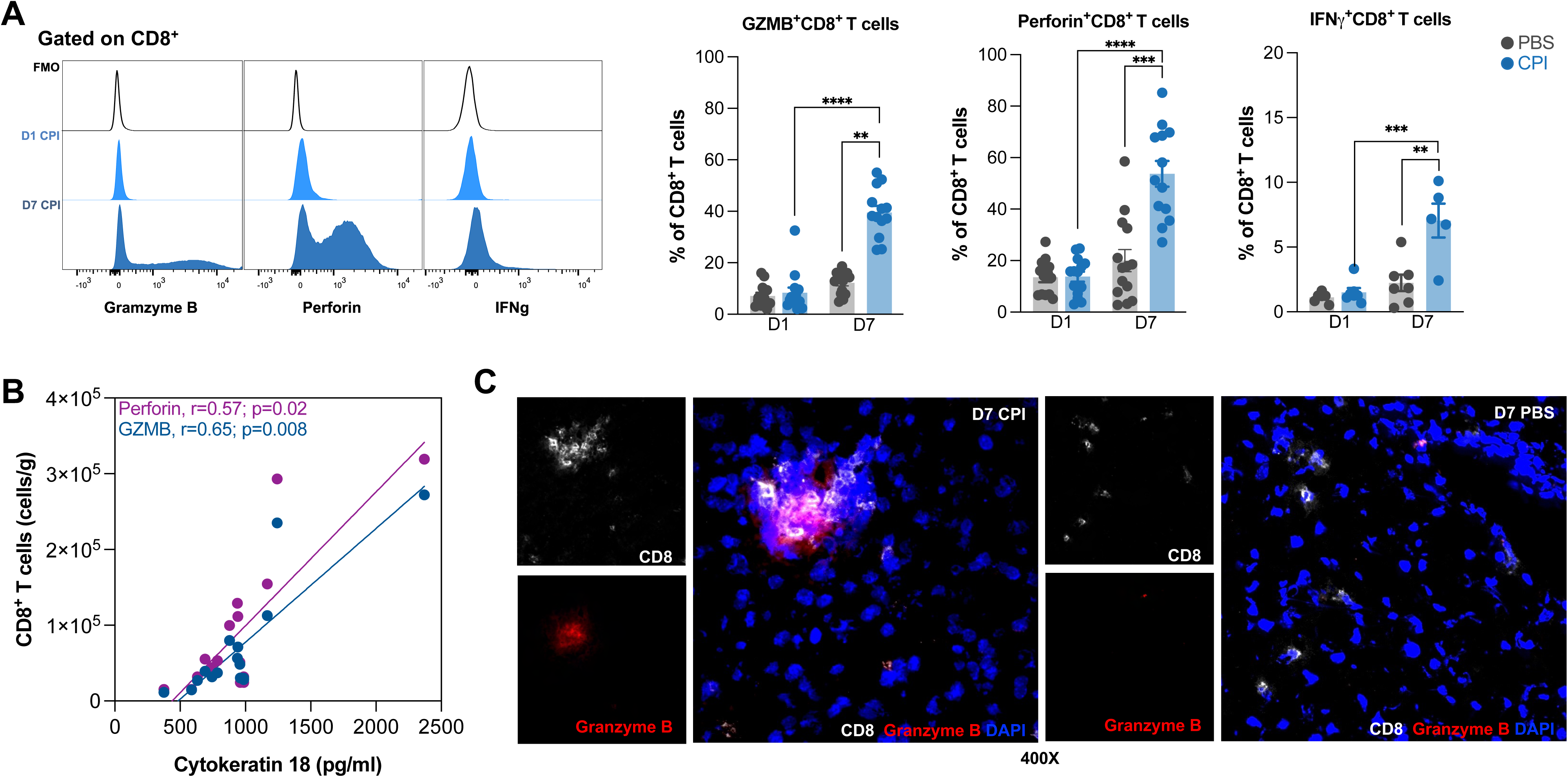
Enhanced cytotoxic CD8^+^ T cell activity in TLR9-L/CPI-treated mice. A) Representative histograms and frequencies of liver GZMB^+^, Perforin^+^ and Ifnγ^+^CD8^+^ T cells in CPI and PBS-treated mice on D1 compared to D7. B) Spearman correlations of absolute numbers of GZMB^+^ (blue) and Perforin^+^ (purple) CD8^+^ T cells with CK-18 levels in D7 CPI-treated mice. C) Representative pictures of liver cryosections stained for CD8 (white), GZMB (red) and DAPI (blue) on D7 of CPI and PBS-treated mice (n=4/group). Magnification: 400X. Each symbol represents an individual mouse. Results shown are representative of 1-3 independent experiments. *p<0.05, ** p<0.01, *** p<0.001. **** p<0.0001.

To identify topographically where myeloid cells accumulate in liver tissue, cryosections were stained for F4/80 and CD11b. CD11b^+^ cells formed lobular inflammatory clusters on D7 in CPI-treated mice (**Supplementary Fig.1 E**, **Supplementary Fig.3 C**), whereas D7 PBS-treated mice showed evenly distributed CD11b^+^ cells. In contrast, F4/80^+^ cells were distributed evenly across lobules and portal tracts in D7 CPI-treated mice with typical KC morphology (**Supplementary Fig.3 C**). Furthermore, mirroring the human liver histology, CD11b/CD8 and F4/80-CD8 double staining revealed that clusters of CD11b^+^ also contained CD8^+^ T cells (**Fig.1 F**, **Supplementary Fig.1 E**), while F4/80 cells were distributed across hepatic lobules and not localised to the same areas (**Supplementary Fig.3 D**). These data suggest that recruited CD11b^+^ monocytes, rather than F4/80^+^ KC, contribute to the formation of inflammatory foci in CPI-hepatitis.

In line with these findings and similarly to human CPI-hepatitis, increased numbers of monocytes expressing the recruitment chemokine receptor CCR2, were measured on D7 following CPI treatment and CCR2 was expressed within inflammatory clusters (**Fig.1 G**, **Supplementary Fig.1 E**). Notably, monocytes are the main expressors of CCR2 within the liver out of other myeloid cells (**Supplementary Fig.3 E**). This liver homing phenotype of monocytes was reflected in significantly increased total liver mRNA *Ccl2* levels (ligand for CCR2) on D7 in CPI-treated mice compared to D7 PBS (**Fig.1 H**). Notably, intracellular cytokine staining revealed that KCs were a potential source of CCL2 (**Supplementary Fig.3 F**).

### Liver transcriptome demonstrates exaggerated and perpetuated inflammatory responses in TLR9-L/CPI treated mice

Our previous human transcriptional analysis of circulating monocytes and CD8^+^ T cells revealed the activation of transcription factors such as *JUN, GATA3, NF_κ_B* and *JAK* signalling, and inflammation associated markers (e.g., *TNFα* and *IL-6*) (18). To gain a deeper insight into the inflammatory environment of the liver throughout the CPI-hepatitis time course, bulk RNA sequencing was performed. Principal component analysis (PCA) and clustered heatmap of differentially expressed (DE) genes across all samples demonstrated similar gene expressions on D1 in PBS and CPI-treated mice and on D4 between both groups (**Fig.2 A+B**). D7 CPI-treated mice demonstrated persistent alterations in their transcriptional profile with augmentation of expression of inflammation-associated genes. In contrast, PBS-treated mice demonstrated only a transient shift in gene expression profile on D4, which resolved by D7. The D7 PBS samples showed clustering with D1 PBS both in PCA space and in DE genes (**Fig.2 A+B**). In addition, DoRothEA predicted transcription factor (TF) activation analysis showed a modest activation of various inflammation-associated TFs on D4 in both CPI and PBS-treated mice, whereas the activation of such TFs (e.g., *Jun, Stat1, Nfkb1, Irf9* and *Gata3*) was further enhanced in D7 CPI-treated mice (**Fig.2 C**), replicating aspects of the reported human transcriptional profile (18). In contrast, TF expression patterns had returned to near baseline in D7 PBS-treated mice (**Fig.2 C**).

These data suggested that the observed changes on D4 are largely TLR9-L treatment effects while D7 changes are due to CPI treatment; therefore, our subsequent analyses focused on the comparisons between D7 TLR9-L/CPI and D7 TLR9-L/PBS mice. RNA sequencing confirmed flow cytometry protein expression analyses (*Ly6c2, Ccr2, Cd8a* and *Gzmb*) and increased mRNA levels of *Ccl2* and further showed differential expression of T cell recruiting chemokines (*Cxcl9/10*) and markers of inflammation (e.g. *Ifnγ, Tnf*α) (**Fig.2 D & Supplemental materials/excel spreadsheet**). KEGG pathway analysis of this D7 comparison showed downregulation of pathways involved in metabolism, particularly associated with anti-inflammatory conditions (e.g. tryptophan metabolism, oxidative phosphorylation) and the upregulation of pathways demonstrating a highly activated, pro-inflammatory profile (**Fig.2 E)**. These included pathways important for cell migration (chemokine signalling, leukocyte endothelial migration), cell interactions (e.g. cytokine-cytokine receptor, T cell receptor signalling, MAPK signalling, antigen presentation), inflammation (e.g. NFκB signalling, JAK-STAT signalling) and cytotoxicity (**Fig.2 E & Supplementary Fig.4 B-E**). More specifically, ingenuity pathway analysis (IPA) identified TNF, Ifnγ, IL-1 and CXCL chemokines as key drivers of experimental CPI-hepatitis (**Supplementary Fig.4 A**).

**Figure 4.**
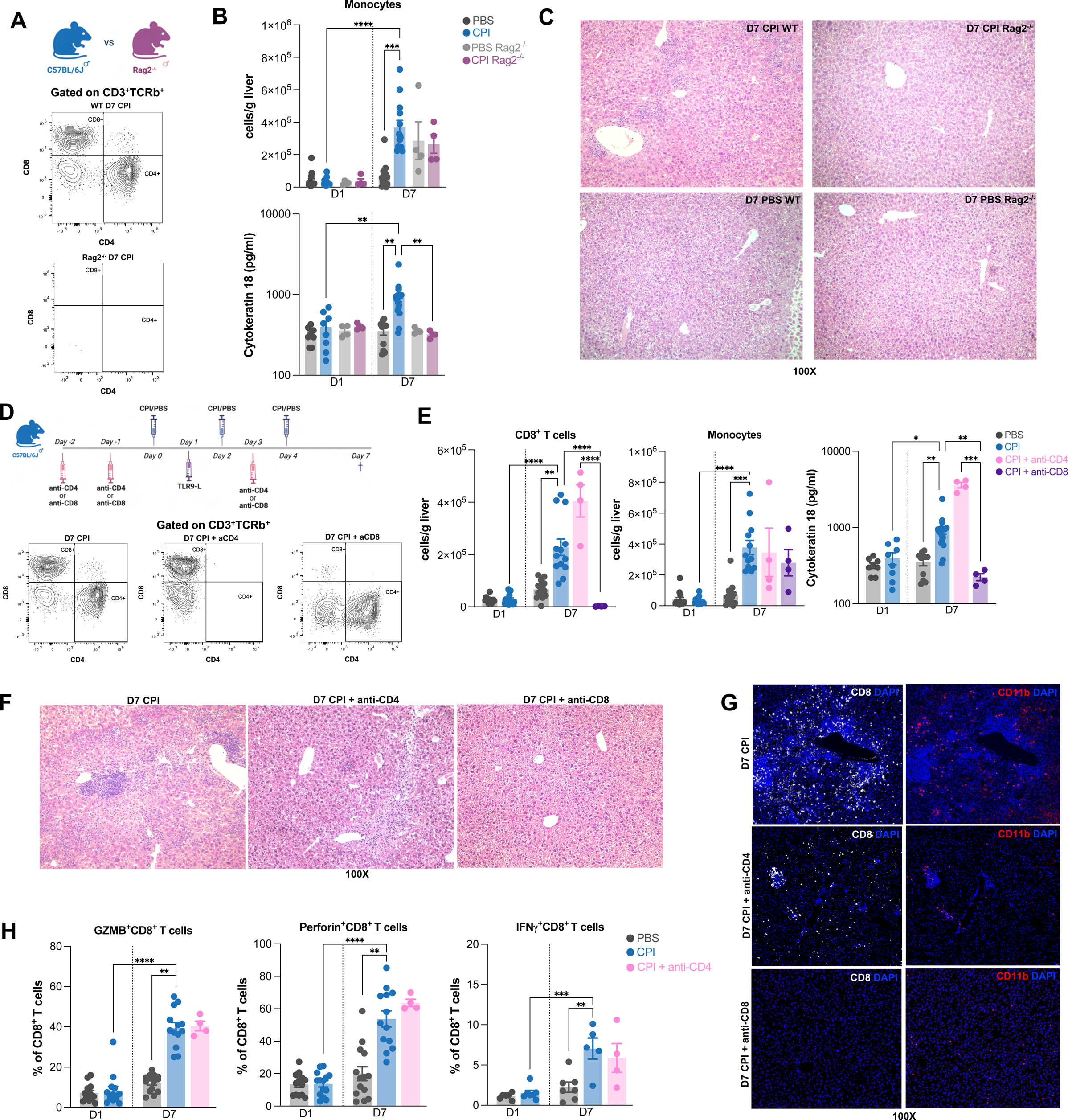
CD8^+^ T cells mediate hepatocyte damage following TLR9-L/CPI treatment. Comparison of experimental CPI-hepatitis time course in WT, Rag2^-/-^ and CD4 or CD8 depleted mice. A) Representative contour plots pre-gated on CD3^+^ confirming the absence of CD8^+^ and CD4^+^ T cells in Rag2^-/-^ mice. B) Absolute numbers of total liver monocytes (top) and measurement of CK-18 plasma levels (bottom) in WT vs Rag2^-/-^ mice on D1 compared to D7. C) Representative H&E stained liver sections of D7 WT vs Rag2^-/-^ mice (n=4/group). Magnification: 100X. D) Depletion schedule of CD4^+^/CD8^+^ T cells during experimental CPI-hepatitis in WT mice. Representative contour plots pre-gated on CD3^+^ confirming the absence of either CD4^+^ or CD8^+^ T cells in WT mice. E) Absolute numbers of CD8^+^ T cells and monocytes and CK-18 plasma levels during the CD4/CD8 depletion time course. F) Representative H&E stained liver sections of mice on D7 following CPI ± anti-CD4/anti-CD8 treatment (n=4/group). Magnification: 100X. G) Representative pictures of D7 liver sections stained left: for CD8 (white) and DAPI (blue) and right: CD11b (red) and DAPI (blue) of D7 CPI ± anti-CD4/anti-CD8 treated mice (n=4/group). Magnification: 100X. H) Frequencies of liver GZMB^+^, Perforin^+^ and Ifnγ^+^CD8^+^ T cells in CPI and PBS-treated mice and D7 CPI+anti-CD4 treated mice. Each symbol represents an individual mouse. Results shown are representative of 1-3 independent experiments. * p<0.05, ** p<0.01, *** p<0.001. **** p<0.0001.

### Increased cytotoxicity of CD8^+^ T cells is associated with hepatocyte damage

Based on reports of enhanced cytotoxicity particularly within inflammatory foci in human CPI-hepatitis and our transcriptional data showing upregulation of genes associated with cytotoxicity, we next assessed the cytotoxic activity of CD8^+^ T cells. Functional analysis showed an expansion of CD8^+^ T cells producing GZMB, perforin and Ifnγ on D7 in CPI-treated mice (**Fig.3 A**). This increase in cytotoxic activity correlated positively with disease severity (GZMB: r=0.65, p=0.008; Perforin: r=0.57, p=0.02) (**Fig.3 B**). Moreover, tissue staining revealed that CD8^+^ T cells produce GZMB within the centre of these immune clusters (**Fig.3 C**).

### CD8^+^ T cells are required for development of CPI-hepatitis

To identify whether liver injury following TLR9-L/CPI treatment is in fact driven by the lymphoid compartment, we performed the CPI-hepatitis time course using Rag2^-/-^ mice (which lack mature lymphocytes) (**Fig.4 A**). While monocyte numbers were similarly increased among groups, liver injury index CK-18 plasma levels were significantly reduced in D7 CPI Rag2^-/-^ mice compared to D7 CPI WT mice (**Fig.4 B**) while Rag2^-/-^ mice also showed normal liver histology (**Fig.4 C**).

To further pinpoint the subset of T cell mediating CPI-hepatitis, we carried out targeted CD4 or CD8 depletion in WT mice during the time course (**Fig.4 D+E**). In the absence of CD4^+^ T cells (D7 CPI + anti-CD4), absolute numbers of CD8^+^ T cells and monocytes were similar to D7 CPI WT mice (**Fig.4 E**). This was reflected by elevated CK-18 levels, histology showing inflammatory foci of CD8^+^ T cells and CD11b^+^ monocytes (**Fig.4 E-G**) and enhanced cytotoxicity of CD8^+^ T cells similar to D7 CPI WT mice (**Fig.4 H**). In contrast, mice without CD8^+^ T cells (D7 CPI + anti-CD8) (**Fig.4 D+E**), despite similarly increased absolute numbers of liver infiltrating monocytes compared to D7 CPI WT mice, were protected from liver injury and showed normal liver histology and distribution of monocytes (**Fig.4 E-G**).

### Monocytes mediate liver recruitment, clustering and cytotoxic activity of CD8^+^ T cells following TLR9-L/CPI treatment

Given our data demonstrating that cytotoxic CD8^+^ T cells drive liver injury in CPI-hepatitis, we next questioned what mediates their liver infiltration. Our transcriptomic data revealed an upregulation of the chemokine receptor *Cxcr3* and its ligands (e.g., *Cxcl9, Cxcl10*) important for T cell recruitment (**Supplementary Fig.4 D**). These findings were confirmed by flow cytometry showing an expansion of CXCR3^+^CD8^+^ T cells on D7 in CPI-treated WT mice (**Fig.5 A**) as well as significantly increased CXCL9/10 plasma levels and total liver mRNA expression (**Fig.5 B**). Cytokine staining identified monocytes as the substantial source for CXCL9 on D7 during experimental CPI-hepatitis (**Fig.5 C**). Notably, the number of liver infiltrating CCR2^+^ monocytes also correlated positively with the number of CD8^+^ T cells producing GZMB and perforin (GZMB: r=0.74, p=0.0015; Perforin: r=0.79, p=0.0005) (**Fig.5 D**).

**Figure 5.**
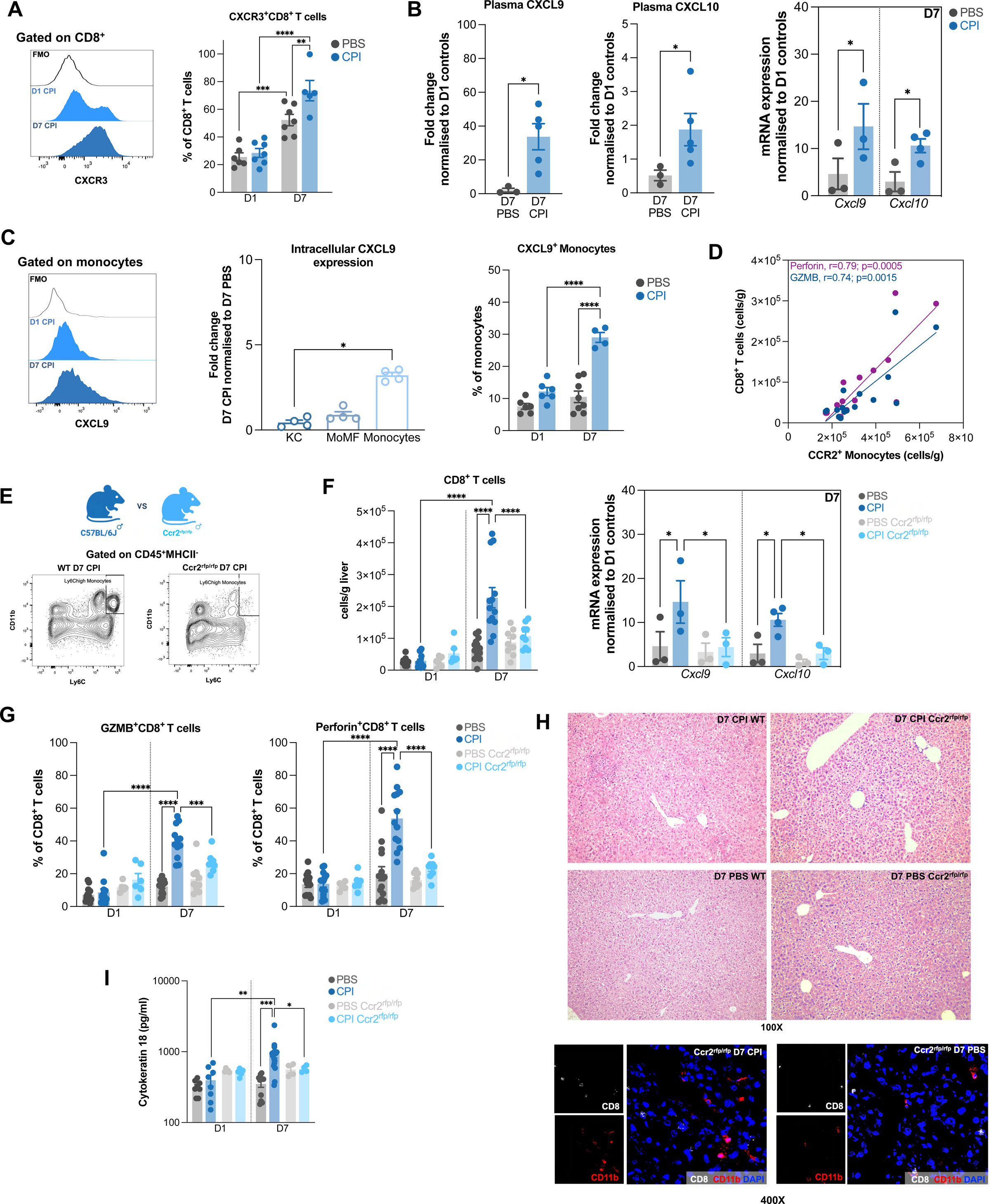
Monocytes mediate recruitment, clustering and activation of tissue destructive CD8^+^ T cells following TLR9-L/CPI treatment. A) Representative histograms and frequency of liver CXCR3^+^CD8^+^ T cells in CPI and PBS-treated mice on D1 compared to D7. B) Fold change plasma CXCL9 and CXCL10 levels (left) and fold change expression of total mRNA levels of *Cxcl9* and *Cxcl10* in liver homogenates (right) in D7 CPI and PBS-treated mice compared to respective D1 controls (n=3-5/group). C) Representative histograms and fold change of CXCL9^+^ liver myeloid cell frequencies on D7 CPI normalised to PBS (n=4/group). Frequency of CXCL9^+^ monocytes in CPI and PBS-treated mice on D1 compared to D7. D) Spearman correlations of absolute numbers of GZMB^+^ (blue) and Perforin^+^ (purple) CD8^+^ T cells with total number of CCR2^+^ monocytes in D7 CPI-treated mice. E) Comparison of experimental CPI-hepatitis time course in WT and Ccr2^rfp/rfp^ mice. Representative contour plots pre-gated on MHCII^-^ cells showing few Ly6C^high^ monocytes in Ccr2^rfp/rfp^ mice, compared to WT mice. F) Absolute numbers of total liver CD8^+^ T cells (left) and fold change expression of total mRNA levels of *Cxcl9* and *Cxcl10* in liver homogenates in D7 CPI and PBS-treated WT and Ccr2^rfp/rfp^ mice compared to respective D1 controls (n=3) (right). G) Data showing frequencies of GZMB^+^ and Perforin^+^CD8^+^ T cells during the time course. H) Representative H&E stained liver sections of D7 WT vs Ccr2^rfp/rfp^ mice during the time course (n=4/group; magnification: 100X) and liver cryosections on D7 stained for CD8 (white), CD11b (red) and DAPI (blue) (n=4/group; magnification: 400X). I) Measurement of CK-18 in plasma. Results shown are representative of 1-3 independent experiments. * p<0.05, ** p<0.01, *** p<0.001. **** p<0.0001.

Since these data, together with observations of dense clusters of CD8^+^ T cells with monocytes, suggest that monocytes may play a role in CD8^+^ T cell recruitment, we next explored the effects of CCR2^+^ monocyte deficiency using Ccr2^rfp/rfp^ transgenic mice (**Fig.5 E**). In Ccr2^rfp/rfp^ mice, the numbers of liver CD8^+^ T cells on D7 did not increase from D1 following TLR9-L/CPI treatment and were significantly reduced compared to D7 CPI-treated WT mice (**Fig.5 F**). This was accompanied by significantly lower levels of *Cxcl9* and *Cxcl10* mRNA (**Fig.5 F**).

The absence of liver infiltrating monocytes, further led to reduced frequencies of CD8^+^ T cells producing GZMB and perforin on D7, compared to CPI-treated WT mice (**Fig.5 G**). Moreover, these mice presented normal liver histology, with evenly distributed CD11b^+^ and CD8^+^ cells (**Fig.5 H**) and baseline CK-18 levels (**Fig.5 I**).

### Therapeutic inhibition of monocyte liver recruitment prevents the development of CPI-hepatitis

Based on our data indicating that CCR2^+^ monocytes are required for CD8^+^ T cell liver recruitment, clustering and activation, we explored the therapeutic potential of monocyte inhibition in preventing CPI-hepatitis development using CVC (a small molecule inhibitor against CCR2/CCR5) in drinking water (**Fig.6 A+B**) (36). Following CVC treatment, significantly fewer CCR2^+^ monocytes were measured on D7 in CPI-treated mice, compared to those receiving vehicle control or normal drinking water (**Fig.6 B**). In line with our previous results, in the absence of CCR2^+^ monocytes, CD8^+^ T cell numbers were significantly reduced on D7, compared to vehicle control or normal drinking water, and were not altered compared to D1 CPI mice (**Fig.6 C**). This was reflected in the reduced frequency of CXCR3^+^CD8^+^ T cells, as well as GZMB and perforin producing CD8^+^ T cells in CVC-treated mice (**Fig.6 D**). The loss of CCR2^+^ monocytes and the associated reduction in CD8^+^ T cell recruitment and cytotoxic activity further resulted in the prevention of the development of liver injury in CVC-treated mice (**Fig.6 E**).

**Figure 6.**
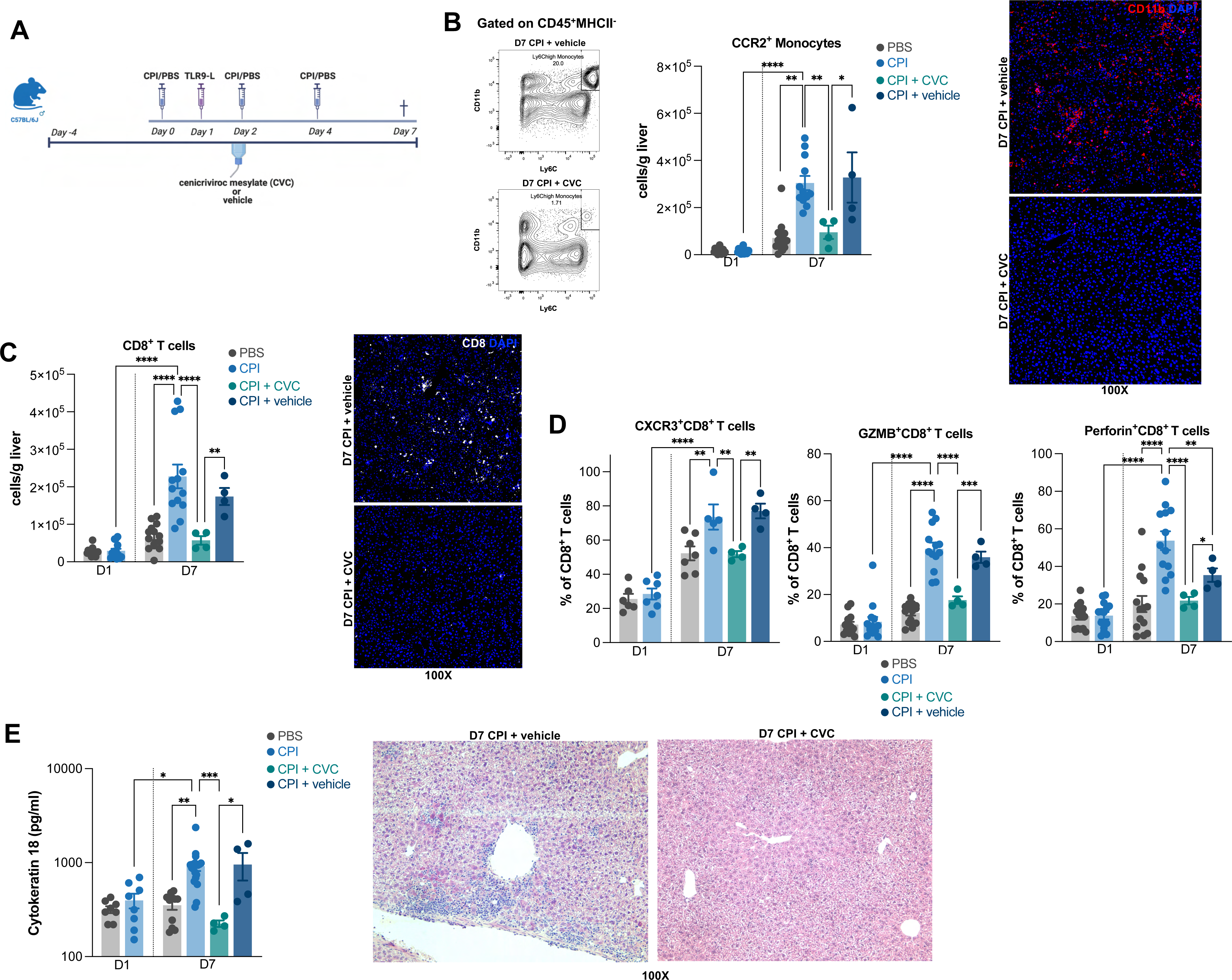
Therapeutic inhibition of monocyte liver recruitment is protective of experimental CPI-hepatitis. A) Set-up of experimental CPI-hepatitis time course and CVC/vehicle treatment schedule. B) Representative contour plots pre-gated on MHCII^-^ cells showing few monocytes in CVC, compared to vehicle treated mice and absolute numbers of total liver CCR2^+^ monocytes during the time course. Representative liver sections of D7 vehicle vs CVC-treated mice stained for CD11b (red) and DAPI (blue) (n=4/group; magnification: 400X). C) Absolute numbers of total liver CD8^+^ T cells (left) and representative liver sections of D7 vehicle vs CVC-treated mice stained for CD8 (white) and DAPI (blue) (n=4/group; magnification: 400X). D) Frequencies of total liver CXCR3^+^, GZMB^+^ and perforin^+^CD8^+^ T cells during the time course. E) Measurement of CK-18 in plasma (left) and representative H&E stained liver sections (n=4/group; magnification: 100X). Results shown are representative of 1-3 independent experiments. * p<0.05, ** p<0.01, *** p<0.001. **** p<0.0001. GZMB: Granzyme B.

## Discussion

In this work, we established a new mouse model of CPI-hepatitis. In line with previous studies, durable hepatitis failed to be induced in the context of CPI alone (29,30). Recently, Hutchinson et al., proposed an association of cytomegalovirus (CMV) reactivation with the development or worsening of CPI-hepatitis, suggesting that the triggered T cell-mediated immunity in the context of CPI might drive liver inflammation directly or promote tissue-damaging bystander T cell responses (42). In our study, administration of CPIs in combination with a TLR9 stimulating agent replicating a viral/bacterial trigger induced a model closely resembling human CPI-hepatitis, thus providing a new platform for further *in vivo* mechanistic studies. Parallel to human liver pathology and circulating immune cell phenotype, mice in our model presented with mixed cytotoxic GZMB^+^perforin^+^CD8^+^ T cell and CCR2^+^ monocyte inflammatory foci with surrounding hepatocyte apoptosis and elevated plasma markers of liver injury (18). Liver RNA sequencing replicated many aspects of the human transcriptomic profile of circulating monocytes/CD8^+^ T cells and identified an exaggerated tissue-damaging inflammatory response characterised by increased expression of key pro-inflammatory mediators (*Nf_K_b, Stat1, Tnfα, Ifnγ*) and chemokines (*Cxcl9, Cxcl10* and *Ccl2*). Using this model, we demonstrate that cytotoxic CD8^+^ T cells are drivers of hepatocellular injury, while their recruitment, clustering and activation is mediated by interactions with liver CCR2^+^ monocytes. The absence of hepatic monocyte recruitment following genetic deletion of CCR2 or through its therapeutic inhibition using CVC prevented the development of experimental CPI-hepatitis.

Our data show that administration of a physiologically plausible priming stimulus in the context of checkpoint inhibition generated hepatitis with a similar histological characteristic of human disease. This differs to other published models of CPI-hepatitis utilising *Pdcd1^-/-^* mice, which are known to have increased CD8^+^ T cell liver infiltration and monocyte activation during homeostasis (28,30,33,43) and supports the theory that CPI-hepatitis arises when a subclinical ‘trigger’ occurs in the context of checkpoint blockade. While the mild TLR9-L induced inflammation resolves by day 7 in PBS-treated mice, loss of important liver tolerance mechanisms in the context of checkpoint inhibition allows perpetuation of unchecked inflammatory responses, ultimately causing hepatocellular damage.

Recent murine studies focusing on priming of hepatic immune responses using TLR9-L as an immunotherapeutic strategy to overcome the exhaustion of CD8^+^ T cells, showed that TLR9-L administration induced local activation and expansion of CD8^+^ T cells within cocoon-like structures of myeloid cells in the liver in the context of chronic viral liver infection (44). Moreover, Cebula et al. showed that the systemic treatment of mice with TLR9-L is beneficial in therapeutic vaccination for immunisation against hepatotropic viruses and the development of memory-recall responses in CD8^+^ T cells (45). TLR9-L administration was also shown to promote the control of tumour growth in murine hepatomas by inducing proliferation of effector CD8^+^ T cells within immune clusters formed with monocytes in the liver (46). Similarly, in experimental CPI-hepatitis, the interaction between CD8^+^ T cells and CCR2^+^ monocytes within inflammatory clusters may lead to the expansion and enhanced cytotoxicity of CD8^+^ T cells.

Using Rag2^-/-^ mice, as well as selective depletion of CD4^+^ and CD8^+^ T cell populations, we reveal that CD8^+^ T cells are main drivers of this damage. This is further supported by recently published data by Llewellyn et al., demonstrating that selective CD8^+^ T cell depletion abrogated hepatotoxicity in another model of CPI-hepatitis (33). Moreover, our analysis identified cytolytic mediators (e.g., GZMB, perforin), Ifnγ and CXCL chemokines as key drivers of experimental CPI-hepatitis, providing additional support for a CD8-mediated mechanism of cytotoxicity directed towards hepatocytes. These pathways have been associated with a number of pathologies including other CPI irAEs (47–50). However, direct targeting of CD8^+^ T cells or the identified relevant mediators would significantly compromise the CPI response, as functional CD8^+^ tumour infiltrating lymphocytes are essential in mediating anti-tumour immunity.

RNA sequencing of liver tissue further supports the role for activation of the TNFα pathway in CPI-hepatitis. While anti-TNFα agents have been reported as a therapeutic option for the treatment of CPI-colitis, the implications for CPI-hepatitis are controversial. Infliximab has recently been associated with the development of hepatotoxicity and thus, has been avoided in these patients (51,52).

Our data further suggests a complex chemokine/cytokine-mediated crosstalk leading to the recruitment and activation of the tissue destructive CD8^+^ T cells by monocytes. The inhibition of initial monocyte recruitment to the liver may therefore present an alternative target upstream of crucial anti-tumour functions. Both, previously reported human (18) and our most recent mouse data highlighted an involvement of the CCR2/CCL2 axis in monocyte recruitment during the pathogenesis of CPI-hepatitis. Genetic deletion of CCR2 in our model prevented the CD8-mediated hepatocyte damage, implicating anti-CCR2/CCL2 as a potential therapeutic avenue. In fact, the role of monocytes in promoting CD8^+^ T cell-associated liver injury and the beneficial effects CCR2/CCL2 targeting has recently been described in a model of in CD8-dependent idiosyncratic DILI (IDILI) (29). In the study, Mak et al. reported that anti-CCL2 treatment of mice reduced hepatic monocyte infiltration attenuated liver injury (29,53). Moreover, CCR2/CCL2 signalling has been implicated in various other liver conditions, such as acute liver injury, fibrosis/cirrhosis, and tumour progression in hepatocellular carcinoma (HCC). Experimental and early clinical investigations have demonstrated the safety and effectiveness of the CCR2/CCR5 small molecule inhibitor, CVC, as an anti-fibrotic treatment for non-alcoholic steatohepatitis (NASH) and alcohol-induced steatohepatitis (36,54–59). However, most recent data from a phase 3 clinical trial showed that CVC did not improve or worsen steatohepatitis/fibrosis om NASH (57). Nevertheless, our data demonstrate that targeting the CCR2/CCL2 axis has the potential to treat liver inflammation in CPI-hepatitis.

Moreover, in the tumour microenvironment, CCR2/CCL2 was shown to contribute to tumour growth and metastasis in a number of tumours. Tumours secret CCL2 to attract monocytes/macrophages, which in turn polarise to an ‘M2-like’ phenotype and promote local immunosuppression and the development of metastasis (60–62). In pre-clinical models, CCR2/CCL2 inhibition, including CVC treatment, was shown to impede the growth and metastasis of HCC, colon carcinoma, prostate and breast cancer by regulating the recruitment and function of tumour-associated macrophages (TAMs), ultimately leading to improved survival of mice (60,63–67). Similarly, in patients with breast, cervix, prostate, HCC and colon cancer, elevated CCL2 levels in plasma were shown to be associated with poor prognosis and metastatic disease (60,63,67–69). CCR2 inhibitors are being trialled for the treatment of pancreatic cancer in humans (70,71) and phase 1b/2 clinical trials are currently investigating the beneficial effects of a CCR2/CCR5 inhibitor in combination with Nivolumab in HCC, non-small cell lung cancer, colorectal and pancreatic cancer (ClinicalTrials.gov Identifier: NCT03184870, NCT04123379). In addition to the role of CCR2 in the promotion of tumours, CCR5 is an important chemokine receptor for regulatory T cell recruitment, which further suppress the tumour microenvironment (72). Thus, CVC or other antagonists, targeting both CCR2 and CCR5, might not only have the potential to attenuate CPI-hepatitis but could also have beneficial effects on the anti-tumour immune response.

One of the limiting factors of this study is that mice in our model do not bear tumours and we cannot predict how the tumour burden would alter hepatic immune responses during experimental CPI-hepatitis. However, our data mirrors human CPI-hepatitis pathology (15,18,37) suggesting that our observations of CPI-hepatitis are likely to be similar in tumour bearing mice. Future work will focus on utilising our model further and assessing the effects of CVC treatment during experimental CPI-hepatitis on a cancer background (e.g., B16 melanoma model). This work will provide important insights into the effectiveness of CVC in treating CPI-hepatitis, but also shed light on its potential to boost anti-tumour immunity. Moreover, mice in our model show mild liver injury compared to other acute liver injury models (e.g., paracetamol-induced liver injury) (73). Human CPI-hepatitis is often characterised as mild disease, which is generally treated with immunosuppressants, but can lead into fulminant hepatic failure and death (15,19,74). Our *in vivo* model should therefore be tuned to investigate different severity levels according to the Common Terminology Criteria for Adverse Events (CTCAE) grading system and can be used to compare effects of immunosuppressive therapies to CVC treatment.

In summary, using our experimental model, we have shown that liver infiltration of CCR2^+^ monocytes is necessary for the development of CD8^+^ T cell driven hepatocyte injury during CPI-hepatitis. Inhibition of CCR2-mediated monocyte recruitment using CVC is a rational target for future investigation. This work provides important insights into how CPI-hepatitis develops and highlights potential therapeutic pathways to treat liver inflammation in this condition.

### Funding

This project was supported by the Royal Marsden Cancer Charity, Rosetrees Trust (A1783, M439-F1 and CF2\100002), Academy of Medical Sciences (Clinical Lecturer Starter Grant to LP), Imperial College London (ICRF fellowship award, to CG and ET), UK National Institute for Health Research (NIHR) and NIHR Imperial Biomedical Research Centre (BRC) and the Imperial College Wellcome Trust Strategic Fund.

## Acknowledgements

BioRender.com was used to create figures of experimental set-up.

## Disclosures

The authors declare that the research was conducted in the absence of any commercial or financial relationships that could be construed as a potential conflict of interest. KJW is now an employee for AstraZeneca (BioPharmaceuticals R&D, Cambridge, UK). No funding or support was received from AstraZeneca.

## Author Contributions

Conceptualisation – CG, CA, LP, ET; Investigation, Data curation, Formal analysis – CG, SA, EM, MM, ET; Interpretation of data – CG, SA, RG, NP, KW, LP, ET; Writing - original draft – CG; Writing - critical revision of manuscript – CG, SA, MT, NP, KW, LP, ET; Funding acquisition – CG, MT, ST, JL, CA, LP, ET

## Abbreviations

ALT: alanine transaminase
CK-18: cytokeratin-18
CPI: checkpoint inhibitor
CPI-hepatitis: checkpoint inhibitor-induced hepatitis
CTLA-4: cytotoxic T lymphocyte antigen-4
CVC: cenicriviroc
DILI: drug-induced liver injury
ELISA: enzyme-linked immunosorbent assay
FFPE: formalin-fixed paraffin-embedded
FMO: fluorescence minus one
GZMB: granzyme B
HBV: hepatitis B virus
HCC: hepatocellular carcinoma
IPA: ingenuity pathway analysis
irAE: immune-related adverse event
KC: Kupffer cells
MoMF: Monocyte-derived macrophages
NK cells: natural killer cells
PCA: principal component analysis
PD-1: programmed cell death-1
PD-L1: programmed cell death-ligand-1
RT-qPCR: quantitative reverse transcription polymerase chain reaction
TF: transcription factor
TLR-L: Toll-like receptor agonist
Tregs: regulatory T cells.

## Supplementary figure legends

**Supplementary figure 1.**
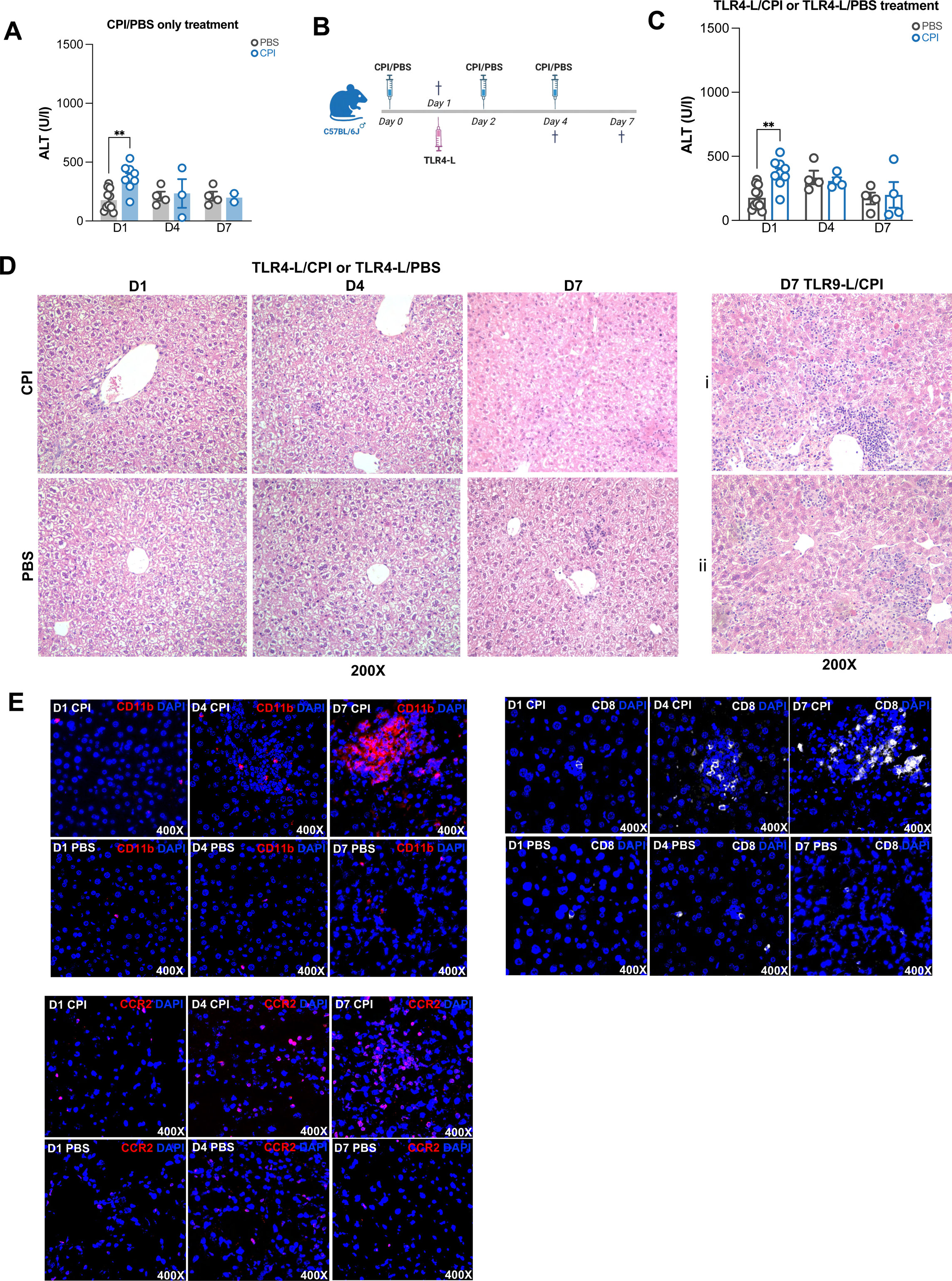
Determination of liver inflammation following CPI only or TLR4-L/CPI treatment of mice. A) Measurement of ALT plasma levels in CPI or PBS only treated mice. B) Experimental set-up of time course comparing CPI and PBS-treated mice across different time points and following TLR4-L administration. C) Measurement of ALT levels in plasma. D) Representative H&E stained liver sections of CPI or PBS-treated mice during TLR4-L time course (n=4/group/time point) compared to D7 TLR9-L/CPI (n=8). Magnification: 200X. E) Representative pictures of liver cryosections stained for CD11b (red) and DAPI (blue); CD8 (white) and DAPI (blue); and CCR2 (red) and DAPI (blue) on D1, D4 and D7 of CPI and PBS-treated mice (n=4/group). Magnification: 400X. Each symbol represents an individual mouse. * p<0.05, ** p<0.01.

**Supplementary figure 2.**
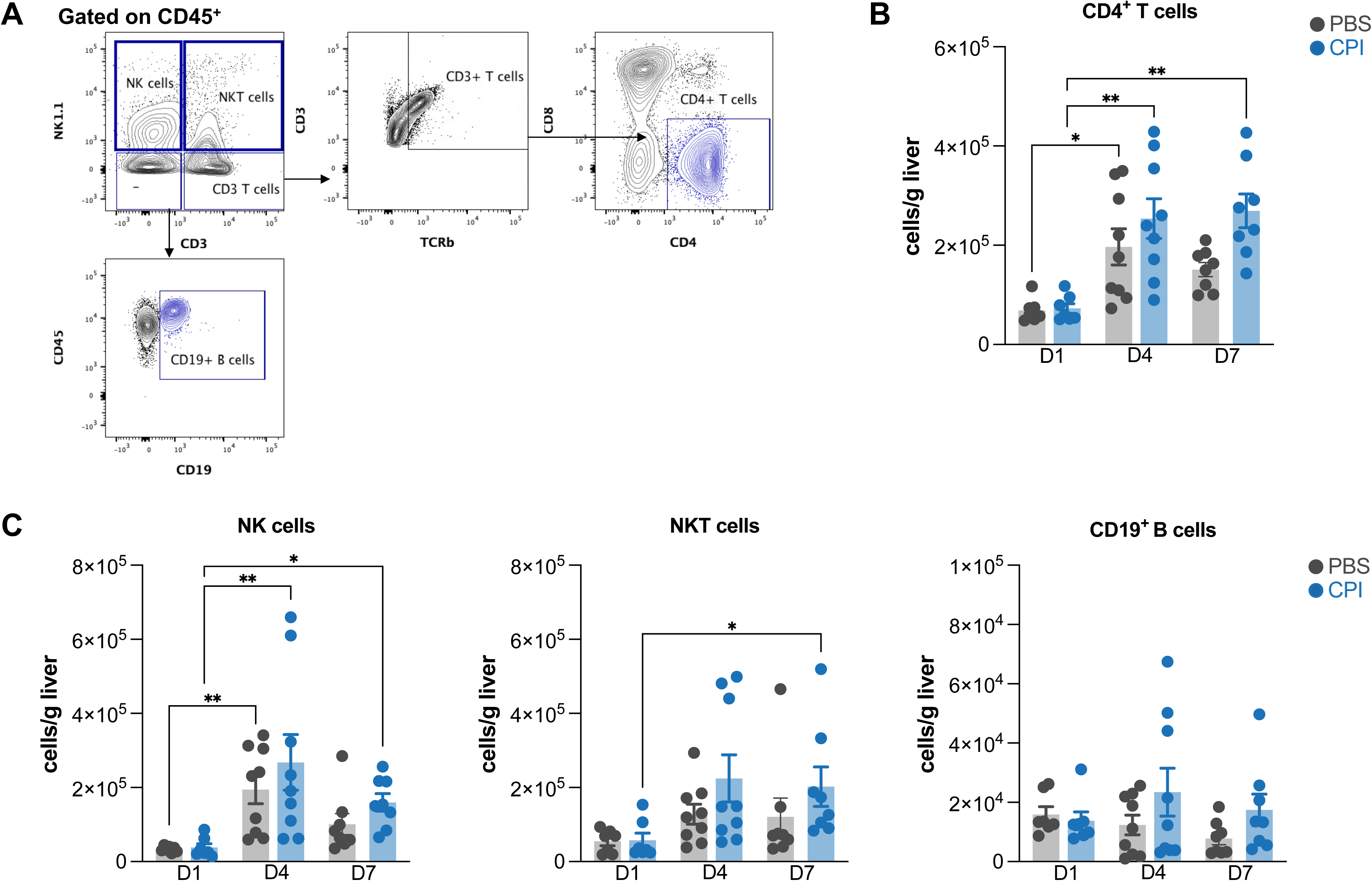
Absolute number of liver lymphocytes following TLR9- L/CPI treatment. A) Representative contour plots of the gating strategy to identify liver lymphocytes. Absolute numbers of CD4^+^ T cells (B) and NK, NKT and B cells (C). Each symbol represents an individual mouse. Results shown are representative of 1- 3 independent experiments. * p<0.05, ** p<0.01.

**Supplementary figure 3.**
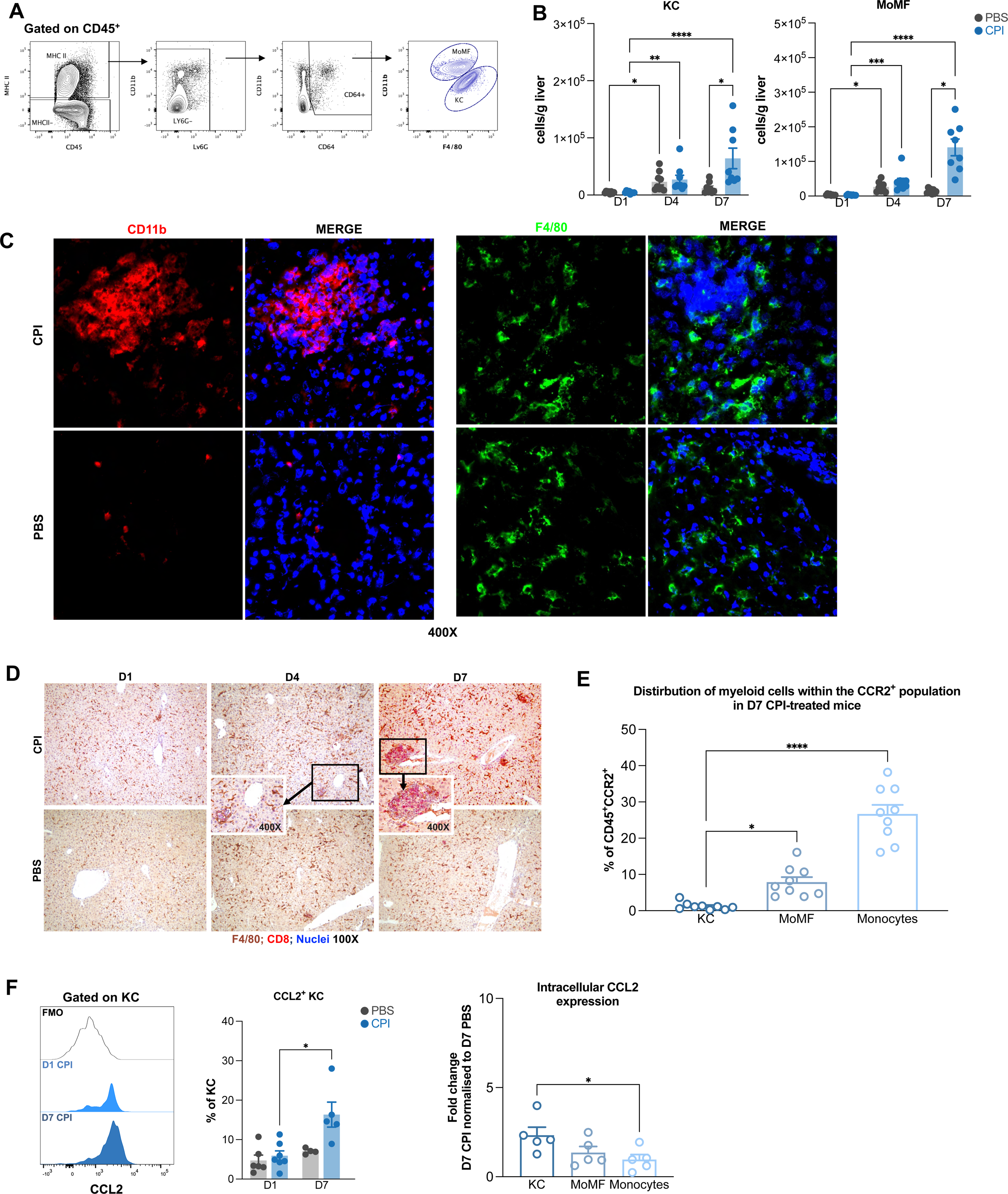
Macrophage distribution in livers following TLR9-L/CPI treatment. A) Representative contour plots of the gating strategy to identify liver myeloid cells. B) Absolute numbers of liver Kupffer cells (KC) and monocyte-derived macrophages (MoMF) in CPI and PBS-treated mice. C) Representative pictures of liver cryosections stained for CD11b (red) and DAPI (blue) (left) and F4/80 (green) and DAPI (blue) (right) on D7 of CPI and PBS-treated mice (n=4/group). Magnification: 400X. D) Representative pictures of FFPE liver sections of D7 CPI or PBS-treated mice stained for F4/80 (DAB, brown), CD8 (Permanent Red, red) and haematoxylin to identify nuclei (n=4/group). Magnification: 100X, 400X. E) Proportions of different myeloid cell populations within CD45^+^CCR2^+^ cells on D7 in CPI-treated mice. F) Representative histograms and frequency of CCL2^+^ KCs in CPI and PBS-treated mice during the time course. Data showing fold change of CCL2^+^ liver myeloid cells frequencies of D7 CPI normalised to D7 PBS (n=4/group). Each symbol represents an individual mouse. Results shown are representative of 1-3 independent experiments. * p<0.05, ** p<0.01, *** p<0.001. **** p<0.0001.

**Supplementary figure 4.**
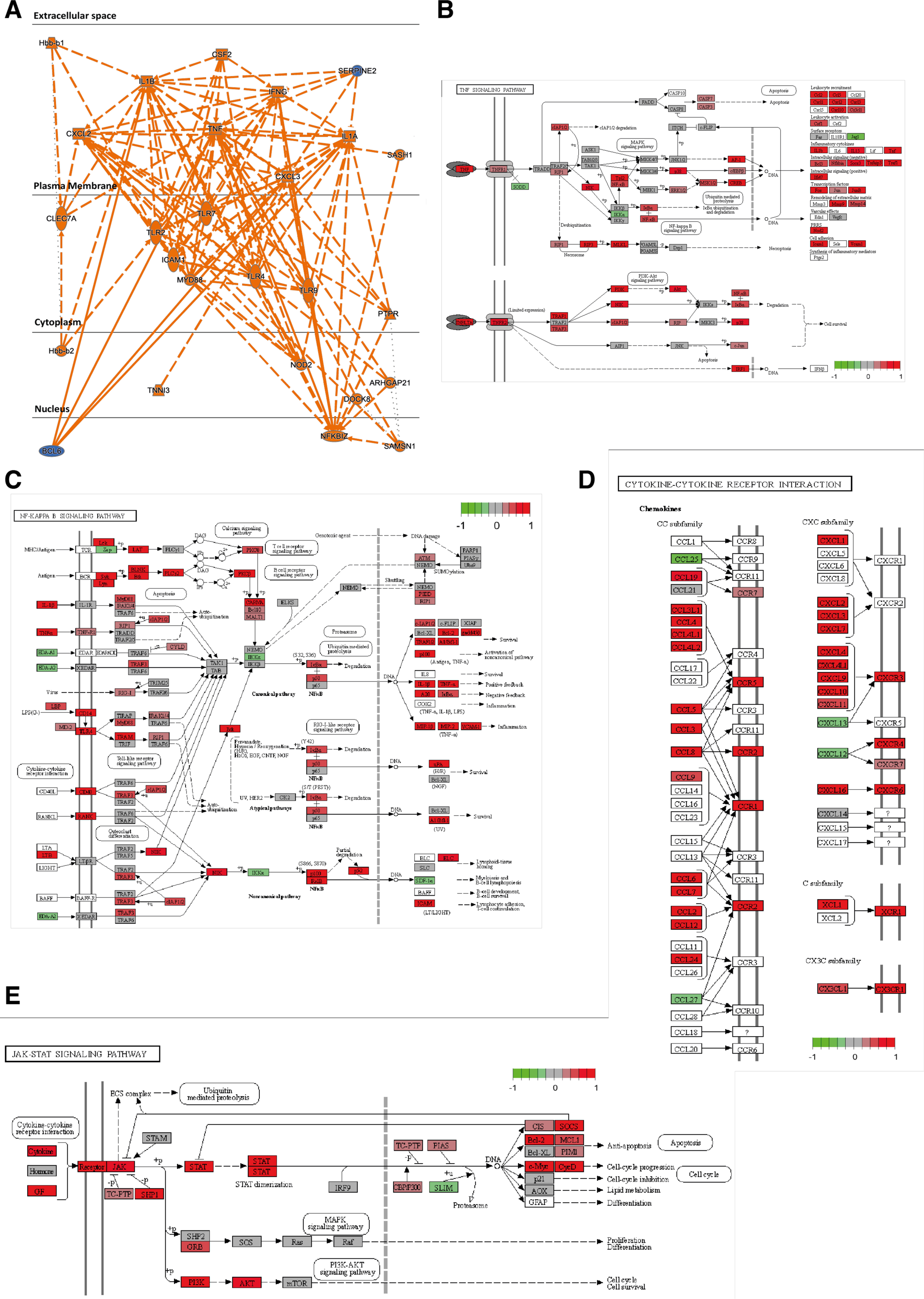
Liver transcriptome predicts TNFα, IL1 and CXC chemokines as upstream drivers of CPI- hepatitis. A) Ingenuity pathway analysis to identify drivers and mediators of CPI-hepatitis. Transcriptomic data mapped to the B) TNFα, C) NF-κB, D) cytokine-cytokine receptor and E) JAK-STAT signalling pathways from KEGG using pathview.

## Notes

### Competing Interest Statement

The authors have declared no competing interest.

