## Supplementary methods for "Therapeutic inhibition of monocyte recruitment prevents checkpoint inhibitor-induced hepatitis"

### Supplemental material and methods

#### Mice and drug treatment

All experimental procedures were carried out in accordance with the UK laws, following the three Rs and with approval of the Home Office and local ethics committees (PPL P8999BD42 and PPL 7007578) in an unblinded manner. C57BL/6 wild-type (WT) mice were used to establish the experimental model of CPI-Hep. The role of lymphocytes in mediating hepatocyte damage was assessed in Rag2$^{-/-}$, lacking all mature lymphocytes, and mice treated with anti-CD4 or anti-CD8 depletion antibodies (all BioXCell). The contribution of monocytes to this pathology was assessed in Ccr2 knockout mice expressing red fluorescent protein (Ccr2$^{rfp/rfp}$) and mice treated with Cenicriviroc (CVC) (MedchemExpress, USA). CVC was dissolved in vehicle containing 0.5% methylcellulose (400cps) (VWR International Ltd, UK) + 1% Tween-80 (Fisher Scientific Ltd, UK) in water and administered in drinking water at 100mg/kg/day.

Stimuli of hepatic inflammation were trialled for the induction of experimental CPI-Hep by the i.p. administration of 20 μg/mouse TLR4 ligand (TLR4-L) [Monophosphoryl Lipid A (MPLA) VacciGrade, InvivoGen] or 20 μg/mouse TLR9 agonist (TLR9-L) CpG oligodeoxynuleotide 1668: 5-S-TCCATGACGTTC CTGATGCT-3) (TIB Molbiol, Germany) on D1, one day post the first administration of CPI or PBS. D1 mice served as baseline control, as these mice did not receive TLR4-L or TLR9-L for hepatic priming.

#### Blood and liver tissue sampling

Mice were sacrificed by terminal anaesthesia receiving 0.2 ml of Pentoject (Centaur Services, UK) i.p. on D1, D4 and D7 post administration of CPIs or PBS. Deep anaesthesia was confirmed by checking paddle and eye reflexes and blood was subsequently collected by cardiac puncture of the right ventricle in blood collection tubes (Microvette, Sarstedt, Germany) to prevent clotting. Mice were then perfused using PBS. Following perfusion, liver tissue was excised and fixed in 10% formalin (Sigma-Aldrich, USA) or Optimal cutting temperature compound (OCT compound; VWR, USA) for histological examination, snap frozen in liquid nitrogen for mRNA analysis or kept on ice cold PBS for fresh cell staining. Plasma was collected by centrifugation of collected blood from the right ventricle.

#### Histology

4 µm thick liver sections from formalin-fixed paraffin-embedded (FFPE) liver tissue were stained with haematoxylin and eosin (H&E) and provided by the Research Histology Facility, Imperial College London.

#### Immunofluorescence microscopy

OCT fixed liver tissue was cut into 15 µm thick cryosections and stored at -80$^{o}$C. Before staining, slides were defrosted and fixed in ice cold acetone for 10 minutes. Slides were subsequently incubated with PBS and 5% BSA for 45 minutes to block unspecific binding. Anti-mouse CD8, F4/80, CD11b, CCR2, GZMB and albumin fluorescently labelled monoclonal antibodies (**Supplementary Tab.1**) were diluted in PBS, 0.1% Triton X-100 and 5% BSA at optimised concentrations and incubated for 1.5 hours at room temperature in a humidity chamber in the dark. Slides were washed with PBS and 0.1% Triton X-100 three times for 5 minutes and once with PBS for 5 minutes before mounting with fluoroshield with DAPI (Sigma-Aldrich, USA). Slides were then stored in the dark at 4$^{o}$C.

**Supplementary Table 1.** List of monoclonal antibodies for immunofluorescent staining of cryosections.

| Antibody | Fluorochrome | Clone | Working concentration | Cat. Number |
|---|---|---|---|---|
| anti-mouse F4/80 | AF488 | BM8 | 10 µg/ml | BioLegend #123120 |
| anti-mouse CD8 | APC | 53-6.7 | 8 µg/ml | BioLegend #100712 |
| anti-mouse Granzyme B | PE | QA16A02 | 4 µg/ml | BioLegend #372208 |
| anti-mouse CD11b | PE | M1/70 | 4 µg/ml | BD #553311 |
| anti-mouse CCR2 | PE | SA203G11 | 2 µg/ml | BioLegend #150610 |

| | | | | |
|---|---|---|---|---|
| anti-albumin | AF647 | n/a | 10 µg/ml | Invitrogen # A34785 |

#### Immunohistochemistry

Double heat-induced epitope retrieval (HIER) immunohistochemistry (IHC) on FFPE tissue was performed to assess the expression of CD8 and F4/80. Tissue was stained manually overnight at $4^{o}$C in a humidity chamber using anti-mouse monoclonal antibodies (**Supplementary Tab.2**). Antigen retrieval was carried out by HIER using EDTA Tris buffer pH 9. Staining was then performed using the EnVision™ G|2 doublestain system – rabbit/mouse (DAB+/permanent red) (Dako, Agilent Technologies, USA) and visualized with DAB and permanent red according to the manufacturer's instructions. Slides were counterstained with haematoxylin (Agilent Technologies, USA). Images were captures with Leica DM4 B microscope (Leica Camera AG, Germany).

**Supplementary Table 2.** List of monoclonal antibodies for immunohistochemistry staining of FFPE sections

| Antibody | Clone | Working concentration | Incubation | Cat. Number |
|---|---|---|---|---|
| anti-mouse F4/80 | SP115 | 1:100 | overnight at $4^{o}$C | Abcam #ab111101 |
| anti-mouse CD8 | EPR21769 | 1:1000 | overnight at $4^{o}$C | Abcam #ab217344 |

#### Quantitative reverse transcription PCR (RT-qPCR)

Snap frozen liver tissue was thawed and homogenised using the TissueLyser II (Qiagen, Germany). RNA was extracted following manufacturer's instructions using the RNeasy Plus Mini Kit (Qiagen, Germany) and measured by nanodrop, reading the optical density at 260-280 nm. 1 µg of total RNA was reversed transcribed to cDNA using SuperScript IV reverse transcriptase with Random Hexamers (Invitrogen, USA). 100ng of cDNA and the 2xSensiMix SYBR Lo-ROX kit (Bioline, UK) were used for quantification of the genes listed in **Supplementary Tab.3** according to manufacturer's guidelines. The abundance of *Gapdh* mRNA was used for reference.

**Supplementary Table 3.** List of primers used for RT-qPCR

| mRNA | Forward | Reverse |
|---|---|---|
| *Cxcl9* | TCGGACTTCACTCCAACACAG | AGGGTTCCTCGAACTCCACA |
| *Cxcl10* | TCTGAGTGGGACTCAAGGGAT | AGGCTCGCAGGGATGATTTC |
| *Ccl2* | CACTCACCTGCTGCTACTCA | GCTTGGTGACAAAAACTACAGC |
| *Gapdh* | CATCACTGCCACCCAGAAGACTG | ATGCCAGTGAGCTTCCCGTTCAG |

#### RNA sequencing

RNA was extracted as described for RT-qPCR. Extracted RNA was checked for sufficient quantity (Nanodrop A280, ThermoFisher, Wilmington, USA) and quality (Bioanalyzer 2100, Agilent, Santa Clara, USA). Library preparation was performed using NEBNext Ultra RNA Library Prep Kit for Illumina (New England BioLabs, Ipswich, USA). Briefly, mRNA was purified from total RNA using poly-T oligo-attached magnetic beads. After fragmentation, the first strand cDNA was synthesized using random hexamer primers followed by the second strand cDNA synthesis. The library was ready after end repair, A-tailing, adapter ligation, and size selection. After amplification and purification, the insert size of the library was validated on a Bioanalyzer 2100 (Agilent, Santa Clara, USA) and quantified by PCR. Libraries were sequenced on Illumina NovaSeq 6000 S4 flowcell (Illumina, San Diego, SUA) using 150bp paired-end reads to a target sequencing depth of 40 million read pairs per sample.

Sequence data were evaluated for quality control issues using fastqc v0.11.9[1]. No adaptor or quality-based trimming was required. Reads were aligned to reference genome GRCm39[2] and corresponding Ensembl genebuild release 104 using the STAR 2.7.9a aligner[3]. Transcript quantification was performed using the RSEM 1.3.3 algorithm[4]. Raw count data were analysed in R v4.0.4 (R Statisical Foundation, Vienna, Austria). Differential expression analyses were conducted using the limma-voom pipeline (limma[5] v3.46.0). Lowly expressed genes were filtered; count data were normalised and transformed and associated precision weights generated using the voom() function. Count data were modelled using a design model incorporating experimental group, blocked on batch and without an intercept (~0+Group+Batch); differential gene expression was estimated for contrasts of interest. A Benjamini-Hochberg false discovery rate correction was applied to resulting p-values. Transcription factor activity was computationally inferred using the dorothea[6] (v1.2.2) package; analyses were restricted to high quality regulons (A, B). Pathway analysis was conducted using Generally Applicable Gene-set Enrichment for Pathway Analysis (GAGE, gage[7] v2.40.2) and murine KEGG pathways (release 99.1) annotated as "signalling" or "metabolism". A Benjamini-Hochberg false discovery rate correction was applied to resulting p-values. Differential expression results for comparisons of interest were mapped to KEGG pathways using the Pathview package (v1.0[8]). Ingenuity Pathways Analysis (IPA September 2021 release; Qiagen, Hilden, Germany) was performed using a gene list filtered for $q<0.05$ and $|logFC|>2$ and under default parameters.

**Isolation of hepatic mononuclear cells**

Liver tissue was mechanically dissociated using scalpels and passed through a 100 µm cell strainer (BD Biosciences, UK). Subsequently, the cell suspension was centrifuged at 60x*g* for 1 min at room temperature to pellet the hepatocytes. Mononuclear cells were then isolated using Optiprep (Sigma-Aldrich, USA) density gradient, according to manufacturer's instructions. Subsequently, red blood cells were lysed for 1 minute with ACK lysis buffer (Thermo Fisher Scientific, USA), followed by a final washing step in PBS.

**Flow cytometry of liver immune cells and absolute cell counts**

Isolated hepatic mononuclear cells were transferred into FACS tubes, resuspended in 100 µl FACS buffer and incubated with TruStain fcX™ (anti-mouse CD16/32) antibody (BioLegend, UK) for 10 minutes prior staining. Surface staining of cells was carried out in the presence of TruStain fcX™ using fluorochrome-labelled monoclonal antibodies listed in **Supplementary Tab.4**, for 25 minutes at room temperature in the dark. Following incubation, cells were washed in PBS and resuspended in 150 µl

FACS buffer and 50 µl 123count eBeads (Thermo Fisher Scientific, UK). For intracellular staining, following the last wash, cells fixed, permeabilised and stained using the True-Nuclear™ Transcription Factor Buffer Set (BioLegend, UK), following manufacturer's instructions. Subsequently, cells were resuspended in 150 µl FACS buffer and 50 µl 123count eBeads (Thermo Fisher Scientific, UK) for acquisition. Fluorescence minus one (FMO) were used as controls.

**Supplementary Table 4.** List of monoclonal antibodies used for flow cytometry.

| Surface marker | Fluorochrome | Clone | Cat. Number |
|---|---|---|---|
| anti-mouse F4/80 | BV421 | BM8 | BioLegend #123137 |
| anti-mouse IFNγ | BV421 | XMG1.2 | BioLegend #505830 |
| anti-mouse Ly6G | BV605 | 1A8 | BioLegend #127639 |
| anti-mouse CXCR3 | BV605 | S18001A | BioLegend #155915 |
| anti-mouse CD45 | BV650 | 30-F11 | BioLegend #103151 |
| anti-mouse CD11b | BV711 | M1/70 | BioLegend #101242 |
| anti-mouse CD8 | BV711 | 53-6.7 | BioLegend #100759 |
| anti-mouse CD3 | BV785 | 17A2 | BioLegend #100232 |
| anti-mouse CCR2 | FITC | SA203G11 | BioLegend #150608 |
| anti-mouse Granzyme B | FITC | QA16A02 | BioLegend #372206 |
| anti-mouse CD8 | FITC | 53-6.7 | Invitrogen #11-0081-85 |
| anti-mouse Perforin | PE | S16009A | BioLegend #154306 |
| anti-mouse CXCL9 | PE | MIG-2F5.5 | BioLegend #515604 |
| anti-mouse CD64 | PerCP/Cy5.5 | X54-5/7.1 | BioLegend #139308 |
| anti-mouse CD19 | PerCP | 6D5 | BioLegend #115532 |
| anti-mouse Ly6C | PE/Cy7 | HK1.4 | BioLegend #128018 |
| anti-mouse NK1.1 | PE/Cy7 | PK136 | BioLegend #108714 |
| anti-mouse CD4 | AF647 | GK1.5 | BioLegend #100424 |

| | | | |
|---|---|---|---|
| anti-mouse MHC Class II | APC-eFluor 780 | M5/114.15.2 | eBioscience #47-5321-82 |
| anti-mouse TCRβ | APC/Cy7 | H57-597 | BioLegend #109220 |
